# Angiotensin II Induces Abdominal Aortic Branch Aneurysms in *Fibrillin-1^C1041G/+^* Mice

**DOI:** 10.1101/2025.11.26.690689

**Authors:** Michael K. Franklin, Deborah A. Howatt, Jessica J. Moorleghen, Mary B. Sheppard, Yuriko Katsumata, Hisashi Sawada, Hong S. Lu, Alan Daugherty

**Author notes:** **Corresponding authors:** Alan Daugherty, Saha CVRC, BBSRB, Room B243, University of Kentucky 741 S Limestone, Lexington KY 40356-0509, E-mail –, Hong S. Lu, Saha CVRC, BBSRB, Room B249, University of Kentucky 741 S Limestone, Lexington KY 40356-0509 E-mail –.

## Abstract

**Background:** Mice harboring a missense variant (C1041G) of *fibrillin-1* (*Fbn1*) have been used extensively for aortopathy research, but do not mimic all facets of the human disease. The role of increased angiotensin II (AngII) or blood pressure in determining the arterial phenotype of these mice remains incompletely defined. The purpose of this study was to determine whether AngII, either directly or indirectly through increased blood pressure, promotes pathology in the proximal thoracic aorta and beyond.

**Methods:** *Fbn1*^*+/+*^ *and Fbn1*^C1041G/+^ littermates were infused with either AngII or norepinephrine (NE) via subcutaneously implanted osmotic pumps. Micro computed tomography (microCT) was used to visualize vascular pathologies. Maximal arterial dimensions were measured using in situ or microCT images.

**Results:** AngII infusion dramatically augmented aortopathy in *Fbn1*^C1041G/+^ mice. Aortic dissection was visible within 3 days of AngII infusion. Over 50% of male *Fbn1*^C1041G/+^ mice died during AngII infusion, primarily due to aortic rupture in either the thoracic or abdominal regions. Surviving males had increased ascending and suprarenal aortic diameters and developed pathological lesions at the celiac and superior mesenteric branches of the abdominal aorta. Female mice had a much lower incidence of death but had increased ascending aortic and branch diameters. Although NE infusion also increased systolic blood pressure, it did not affect mortality or enlarge aortic or branch diameters in *Fbn1*^C1041G/+^ mice. MicroCT identified novel pathological changes during AngII infusion, including development of aortic branch aneurysms in the celiac and superior mesenteric arteries; however, the maximal diameters of the adjacent suprarenal aorta showed only modest increases in male *Fbn1*^C1041G/+^ mice.

**Conclusion:** AngII exacerbated aortic pathology in *Fbn1*^C1041G/+^ mice. It also promoted development of pathologies at aortic branch points, including the celiac and superior mesenteric arteries.

## INTRODUCTION

Fibrillin-1 (FBN1) is a large (∼350 kDa) protein that is hypothesized to be required for the formation of a microfibrillar scaffold upon which elastic fibers are assembled. Genetic variants of *FBN1* are the underlying cause of Marfan syndrome. The manifestations of Marfan Syndrome are highly diverse, which may be at least partially attributable to the fact that there are over 2,000 variants.^1^ While attention has commonly focused on the aberrant dilation of the aortic root in patients with Marfan syndrome, it is increasingly recognized that these individuals have diverse vascular pathologies. These include aneurysm, dissection, and rupture in aortic areas beyond the root, including the ascending, descending thoracic, and abdominal regions.^2^ Aneurysms also frequently occur at multiple aortic branch points. This includes aortic branches arising from the aortic arch (brachiocephalic, carotid, subclavian), descending thoracic (vertebral, bronchial), suprarenal (celiac, superior mesenteric, renal), and infrarenal regions (iliac).^3^ The presence of ancillary aneurysms further increased the complexity of pathologies induced by *FBN1* genetic variants.

While *FBN1* variants cause a diversity of vascular phenotypes, experimental models using *FBN1* variants are predominantly focused on the proximal thoracic aorta.^4^ The most commonly used model is mice expressing a *Fbn1*^C1041G/+^ variant.^5^ These mice have a modest expansion of the aortic root and ascending aorta,^6, 7^ which exhibits profound sexual dimorphism, with progressive enlargement only occurring in male mice up to one year of age.^7, 8^ There have been no reports of aortic dissection or rupture in *Fbn1* ^C1041G/+^ mice.

To determine the role of angiotensin II (AngII) in arterial diseases, many studies have employed chronic subcutaneous infusions, most commonly at a rate of 1,000 ng/kg/min.^9^ This procedure in wild type mice promotes perivascular fibrosis in many arterial beds.^10^ When combined with hyperlipidemia and BAPN administration, AngII infusion promotes localized pathologies with augmentation of proximal thoracic aortopathy and progressive expansion and rupture of the suprarenal aorta.^11-13^ AngII has been infused into *Fbn1*^C1041G/+^ mice. This includes very high infusion rates of AngII (4.5 mg/kg/d = 3,125 ng/kg/min) that increased ascending aorta diameters, with some studies having all mice die of ascending aortic rupture within 4 weeks of infusion.^14-16^ In addition, AngII infusion at 1,000 ng/kg/min in *Fbn1* ^C1041G/+^ mice in a normolipidemic background^17^ or in an ApoE deficient background resulted in increased aortic aneurysms in both the ascending and abdominal regions.^18^

Despite several recent studies on aortic aneurysms,^14-18^ the role of increased AngII activation in aortic pathology has not been extensively investigated in *Fbn1*^C1041G/+^ mice. Therefore, this study was designed to address the following questions: 1. Whether the sexual dimorphism of vascular diseases that occur in *Fbn1* ^C1041G/+^ mice is retained under AngII activation. 2. Whether pathologies develop beyond the proximal thoracic aorta in a mode that recapitulates the human disease. 3. Whether blood pressure per se augments vascular diseases or whether the effect is specific to AngII activation.

## METHODS

### Data Availability

Detailed materials and methods are available in this manuscript. Numerical data are available in the Supplemental Excel File.

### Mice

Studies were performed in accordance with recommendations for design and reporting of animal aortopathy studies.^19^ All experiments used littermate controls. *Fbn1*^C1041G/+^ (stock #012885) mice were obtained from The Jackson Laboratory as described previously.^8^ Male and female *Fbn1*^C1041G/+^ mice were bred to C57BL/6J female and male mice, respectively, to generate male and female *Fbn1*^+/+^ and *Fbn1*^C1041G/+^ littermates. Littermates were separated by sex and randomly assigned to housing groups after weaning. Mice were housed up to 5 per cage, maintained on a 14-hour light/10-hour dark cycle, fed Teklad irradiated global 18% protein rodent diet (Diet # 2918) ad libitum, and allowed ad libitum access to water via a Lixit system. Bedding was provided by P.J. Murphy (Coarse SaniChip) and changed weekly during the study. Cotton nestlets were provided as enrichment. The room temperature was maintained at 21-23 °C, and the humidity was maintained at ∼ 50%.

Both male and female mice were studied. Animal numbers at study initiation and at termination are detailed in the Major Resources Tables. All experiments were approved by the University of Kentucky IACUC (Protocol # 2018-2967).

### Genotyping

Mice were genotyped after weaning and termination, respectively, using tail tissue: group allocation was based on genotyping performed after weaning at postnatal day 28, and the genotype was confirmed again using tissue acquired at the termination of each study. *Fbn1*^C1041G/+^ was assayed using forward primer (5’-CTCATCATTTTTGGC CAGTTG-3’) and reverse primer (5’-GCACTTGATGCACATTCACA-3’) covering a loxP-flanked neomycin resistance cassette placed in intron 24, which is not present in wild type mice. The protocol used was as described by The Jackson Laboratory. *Fbn1*^+/+^ generates a 164-bp product, and *Fbn1*^C1041G/+^ generates a 212-bp product. Post-termination validation genotyping was performed by Transnetyx in a blinded manner.

### Subcutaneous Infusions

After random assignment, AngII (1,000 ng/kg/min dissolved in saline; Product # 4006473; CAS # 4474-91-3; Bachem) or its vehicle (saline), or norepinephrine (NE 5.6 mg/kg/day; dissolved in saline containing 0.2% wt/vol ascorbic acid^20^) or its vehicle (0.2% wt/vol ascorbic acid in saline) was infused through a subcutaneously implanted osmotic pump (ALZET LLC). Alzet model 2001 was used for 3 days, and model 1004 for 28 days of infusion, respectively.^21^ Surgical staples used to close incision sites were removed 7-10 days after surgery.

### Systolic Blood Pressure Measurements

Systolic blood pressure was measured on conscious mice by a non-invasive tail-cuff system (MC4000 Multi Channel system, Hatteras Instruments) following our standard protocol.^22^ Data were collected at the same time each day for 3 consecutive days. Criteria for accepted data were systolic blood pressure between 70 and 200 mmHg and standard deviation < 30 mmHg for at least 5 successfully recorded data/mouse/day. The mean systolic blood pressure of each mouse from the 3-day measurements was used for data analysis.

### Necropsy

All study mice were checked at least once every day. Necropsies were performed immediately to determine the cause of death after carcasses were found. Aortic rupture was defined as the presence of extravascular blood that accumulated in the body cavity. The location of blood egress was determined by the location of the blood clot and a discernible disruption of the aortic wall.

### Micro Computed Tomography (microCT)

MicroCT was performed as described previously.^23^ Mice were euthanized by an overdose of ketamine and xylazine cocktail (90 and 10 mg/kg, respectively). The thoracic cavity was cut open, and the right atrium was nicked to allow the exit of blood flow. Saline (5 ml) was perfused through the left ventricle. The right atrium was sealed using superglue immediately after perfusion, and Microfil® (Flow Tech, Inc.) or Vascupaint (MediLumine Inc.) was injected through the same catheter. Once Microfil or Vascupaint was visualized in the arterioles surrounding the small intestine, the catheter was clamped shut to prevent backflow of Microfil or Vascupaint into the thoracic cavity, and the animal was set aside to allow the compound to harden for ∼90 minutes.

After Microfil or Vascupaint perfusion, animals were scanned using a Skyscan 1276 MicroCT (Bruker) or a U-CT Optical Imaging system (MILabs, Netherlands). MicroCT-scanned images were reconstructed using the N-Recon program (Bruker, Belgium) or MILabs reconstruction program (MILabs, Netherlands) to adjust for beam hardening and ring artifacts. 3D reconstruction and measurements were performed using the 3D slicer program. All bones and vasculatures not of interest were removed using a scissor tool within the program to display the aorta and its major branches. To visualize branch arteries, all the other surrounding vasculatures were removed using the scissor tool. Maximal diameters of suprarenal aortas and aortic branches, including celiac, superior mesenteric, right renal, and left renal arteries, were measured using microCT 3D images.

### Measurement of in situ Aortic Diameters

Mice were terminated by overdose of a ketamine and xylazine mixture followed by cardiac puncture and saline perfusion. The order in which mice were terminated was randomized. Aortas were dissected away from the surrounding tissue. A black plastic sheet was inserted beneath the aorta and heart to increase contrast and facilitate visualization of aortic borders.^24^ Optimal Cutting Temperature compound (Sakura Finetek) was introduced into the left ventricle to maintain aortic patency before imaging. Aortas were imaged using a Nikon SMZ25 stereoscope, and measurements were recorded using NIS-Elements AR 5.11.03 software (Nikon Instruments Inc.). Maximal ascending aortic diameters were measured at the largest width perpendicular to the vessel. Aortic lengths were measured from the beginning of the left subclavian artery to the iliac bifurcation.

### Pathology

Thoracic aortas with major branches were harvested from surviving mice after 3 days of AngII infusion and immersed in neutrally buffered formalin (10% wt/vol). Aortas were embedded in OCT and sectioned with a cryostat.

Abdominal aortas with major branches were dissected free and immersed in neutrally buffered formalin (10% wt/vol) overnight, followed by a series of dehydration steps in increasing concentrations of ethanol. The aortas were then embedded in paraffin wax. Tissue sections (5 μm) were collected with a microtome. Paraffin-embedded sections (5 µm) were deparaffinized using limonene (Cat # 6533A, Medical Chemical Corporation).

Hematoxylin and eosin (H&E, Cat # 26043-06, Electron Microscopy Sciences; Cat # AB246824, abcam) and Verhoeff iron hematoxylin staining were performed, respectively, to visualize pathologies and elastic fibers. Picrosirius red/methyl green staining was performed to visualize collagen fibers. Immunostaining was performed using primary and secondary antibodies listed in the Supplemental Materials Major Resources Tables. NovaRed (Cat #SK-4805, Vector) was used as chromogen. Images of histological staining and immunostaining were captured using an Axioscan 7 (Zeiss) or Nikon Eclipse N*i* and imaged using ZEN v3.1 blue edition (Zeiss) or NIS-Elements AR 5.11.03 software (Nikon Instruments Inc)

### Statistical Analyses

Statistical analyses were performed using R version 4.5.2. For data presented in Figures 1K, 1L, 2B for right and left renal arteries, Figures S2A, S2B, S2F, S2J, and S6, normality and homogeneous variance assumptions for data were assessed using the Shapiro-Wilk test and Levene’s test, respectively. All data that met the normality assumption also satisfied the homoscedasticity assumption. Therefore, two-way analysis of variance (ANOVA), followed by planned contrast tests based on our research hypotheses, was used to compare group means. For data that did not pass the normality test, an aligned rank transform was applied to conduct a nonparametric two-way ANOVA using the “art” function from the “ARTool” R package (version 0.11.2), followed by contrast testing with the “art.con” function. For systolic blood pressure data presented in Figure S3A, since the data passed the normality but not equal variance test, Welch’s t-test was used. For data presented in Figure S3C (N<6/group), Mann-Whitney Rank Sum test was used. For survival analyses, Kaplan–Meier curves were generated and compared using a global log-rank test to assess overall differences among groups, followed by pairwise log-rank tests for between-group comparisons. Statistical significance was set at Bonferroni-corrected *P<0*.*05* for multiple comparisons.

**Figure 1.**
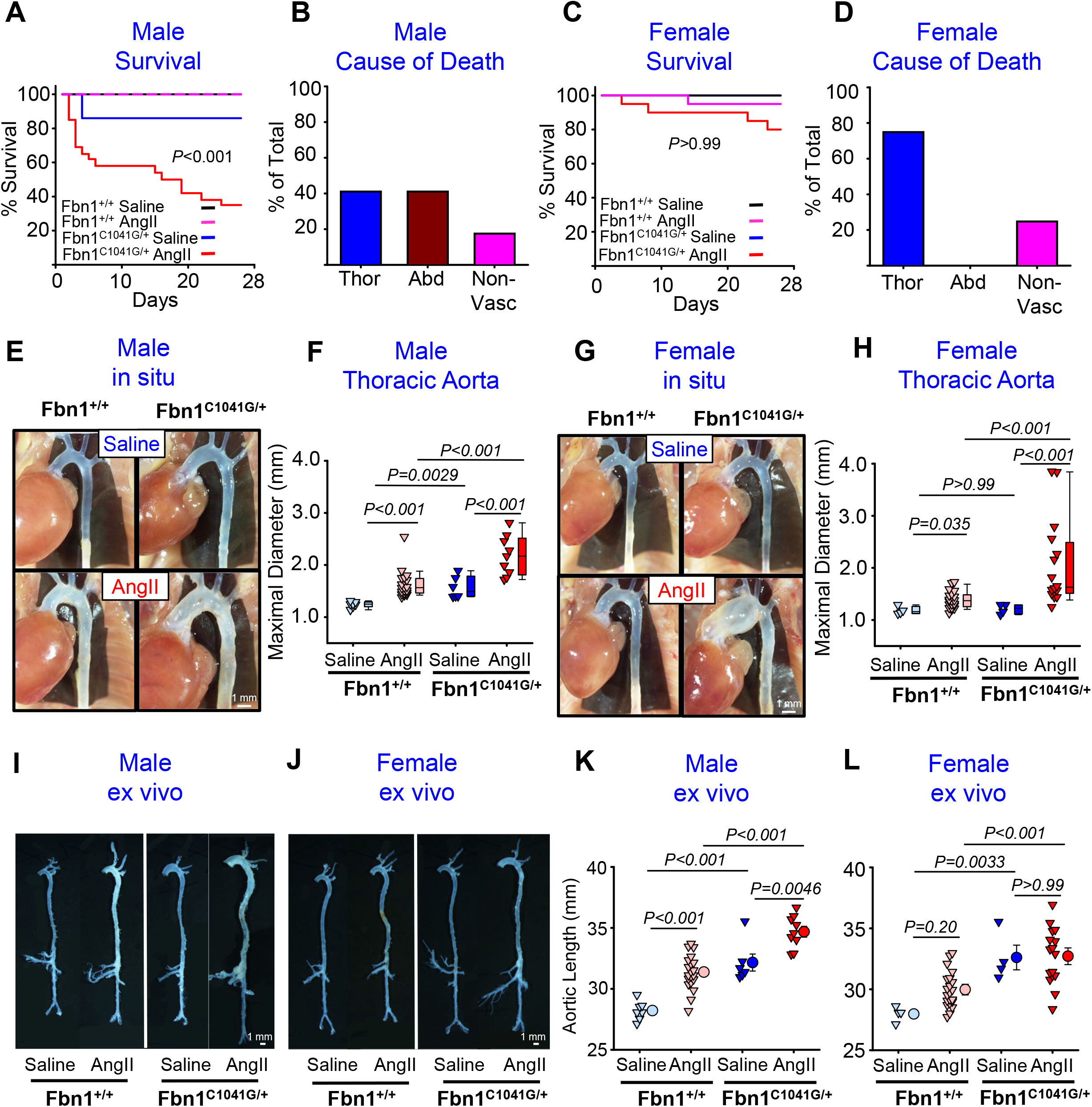
AngII Infusion Augmented Ascending Aortic Aneurysm in *Fbn1*^*C1041G/+*^ mice and Induced Aortic Rupture. Nine-to 14-week-old male and female *Fbn1*^*+/+*^ and *Fbn1*^*C1041G/+*^ mice were infused with either saline or AngII for 28 days. Survival curves and cause of death in male (**A-B**) and female (**C-D**) mice. P values (**A, C**) were determined by Pairwise log-rank analysis. *P*< 0.*001* (**A**) and *P* >0.*99* (**C**) between *Fbn1*^*+/+*^ and *Fbn1*^*C1041G/+*^ mice infused with AngII. Representative in situ images of the thoracic aorta and maximal diameters of ascending aortas in male (**E-F**) and female (**G-H**) mice. Representative ex vivo images of the entire aorta in male (**I**) and female (**J**) mice. Aortic length measured from the left subclavian branch to the iliac bifurcation in male (**K**) and female (**L**) mice. Data were analyzed using two-way ANOVA (nonparametric in **F** and **H** and parametric in **K** and **L**) followed by contrast tests with Bonferroni correction. Male mice: N=7 in *Fbn1*^*+/+*^ saline, N=20 in *Fbn1*^*+/+*^AngII, N=6 in *Fbn1*^*C1041G/+*^ saline, and N=9 in *Fbn1*^*C1041G/+*^ AngII groups at termination. Female mice: N=4 in *Fbn1*^*+/+*^ saline, N=21 in *Fbn1*^*+/+*^AngII, N=4 in *Fbn1*^*C1041G/+*^ saline, and N=16 in *Fbn1*^*C1041G/+*^ AngII groups at termination. Thor: thoracic cavity; Abd: abdominal cavity; Non-Vasc: non-vasculature.

## RESULTS

### AngII Infusion Reduced Survival in Male *Fbn1*^C1041G/+^ Mice

Infusion of AngII for 3 days led to distinctive blood accumulation in both the ascending and descending thoracic aorta of *Fbn1*^C1041G/+^, but not *Fbn1*^+/+^, mice. This phenotype was restricted to the outer margins of the media (Figure S1), as has been demonstrated in several other modes of inducing thoracic aortic pathology in mice.^21, 25^

AngII infusion into male *Fbn1*^C1041G/+^ mice for 28 days led to a high incidence of death (∼65%) that was significantly higher compared to the mortality rate in *Fbn1*^+/+^ mice (Figure 1A; Bonferroni corrected *P<0*.*001*). Necropsies of all mice that died within 14 days of AngII infusion revealed characteristic blood accumulation in either the thoracic or abdominal cavity, with approximately equivalent incidence (Figure 1B). Since the precise loci of blood egress could not be reliably identified, rupture sites were recorded as thoracic or abdominal. All the deaths due to abdominal aortic rupture occurred within the initial 7 days of AngII infusion. Deaths due to thoracic aortic rupture were also prominent within the first 7 days of AngII infusion, but some occurred during more protracted infusion. Some deaths occurred late during AngII infusion that were not discernible as being due to loss of vascular integrity (Supplemental Table 1). In contrast to the high mortality observed in male mice, female *Fbn1*^*C1041G/+*^ mice exhibited a low incidence (20%) of death during AngII infusion, which did not differ significantly from that of *Fbn1*^+/+^ mice (Figure 1C). All the vascular deaths in females were attributed to thoracic aortic rupture (Figure 1D).

Aortas were visualized in mice that survived 28 days of AngII infusion, and maximal diameters of ascending aortas were measured using in situ images, as described previously.^24^ Aortas of both *Fbn1*^+/+^ and *Fbn1*^*C1041G/+*^ male mice infused with saline had a similar opacity, which was increased following AngII infusion (Figure 1E). AngII infusion significantly increased ascending aortic diameters in both *Fbn1*^+/+^ and *Fbn1*^*C1041G/+*^ male mice (Figure 1F). Female *Fbn1*^+/+^ and *Fbn1*^*C1041G/+*^ mice also had similar opacity during saline infusion (Figure 1G). AngII infusion into *Fbn1*^+/+^ female mice failed to promote an overt change in opacity, but ascending aortic diameters were increased modestly. AngII infusion into female *Fbn1*^*C1041G/+*^ mice promoted grossly discernible pathology in the ascending aorta and pronounced increases in diameters (Figure 1H).

Aortas were dissected free to determine whether pathologies were present in any other regions of the aorta. During this tissue processing, longitudinal tortuosity in *Fbn1*^*C1041G/+*^ mice infused with AngII was noted. Therefore, aortic lengths were measured from the left subclavian branch to the iliac bifurcation. AngII infusion significantly increased the length of the aorta, irrespective of *Fbn1* genotype (Figure 1K). The length of the aorta was much greater in male *Fbn1*^*C1041G/+*^ mice and was significantly increased during AngII infusion (Figure 1K). In contrast, the aortic length was not different between saline and AngII infusion in female *Fbn1*^+/+^ and *Fbn1*^*C1041G/+*^ mice, respectively (Figure 1L).

### Norepinephrine Increased Blood Pressure but Failed to Induce Pronounced Aortic Disease

To determine whether increased systolic blood pressure is the primary factor through which AngII infusion augments aortic pathologies in *Fbn1*^*C1041G/+*^ mice, this strain was infused with NE. Infusion of NE at a rate of 5.6 mg/kg/day led to significant increases in systolic blood pressure in both male and female *Fbn1*^+/+^ and *Fbn1*^*C1041G/+*^ mice (Figure S2A and B). In contrast to AngII infusion, there were no deaths in either male or female *Fbn1*^+/+^ or *Fbn1*^*C1041G/+*^ mice during NE infusion (Figure S2C and D). Also, there were minimal changes in the in situ appearance of aortas during NE infusion in either male or female *Fbn1*^+/+^ and *Fbn1*^*C1041G/+*^ mice (Figure S2E and G), nor in the maximum external diameter of the ascending aorta (Figure S2F and H). Furthermore, the increased systolic blood pressure induced by NE infusion failed to alter aortic length or develop any grossly apparent aortic pathologies (Figure S2I-L).

### AngII Promoted Development of Abdominal Aortic Branch Aneurysms in *Fbn1*^*C1041G/+*^ Mice

Ex vivo examination of aortas revealed the development of aortic branch aneurysms in the abdominal region of AngII-infused *Fbn1*^*C1041G/+*^ mice. To gain further insight, aortic imaging was performed using microCT in groups of saline and AngII-infused mice, respectively. No abdominal aortic branch aneurysms were detected in male and female *Fbn1*^+/+^ and *Fbn1*^*C1041G/+*^ mice following infusion with saline, or in *Fbn1*^+/+^ male and female mice infused with AngII (Figure 2A and S3A). In contrast, AngII infusion into *Fbn1*^*C1041G/+*^ male and female mice produced marked aneurysmal expansion in the celiac and superior mesenteric arteries (Figures 2B and S3B). AngII infusion did not induce dilation of the left and right renal arteries in female *Fbn1*^*C1041G/+*^ mice; in male mice, it resulted in a modest increase in the diameters of the left, but not the right, renal artery (Figure 2B and S3B). The increased aneurysm at the aortic branches of the celiac and superior mesenteric arteries was a highly consistent pathology in *Fbn1*^*C1041G/+*^ mice (Figure 2, S3, and Videos 1 and 2). The pathology of aneurysm at both the celiac and superior mesenteric arterial branches was characterized by comparable degrees of elastic fragmentation and regeneration, collagen deposition, expansion of the number of cells expressing α-smooth muscle cell actin, and the presence of numerous CD68 positive cells (Figures 2C and S5), when compared to tissues from the same regions of *Fbn1*^+/+^ and *Fbn1*^*C1041G/+*^ mice infused with saline or *Fbn1*^+/+^ mice infused with AngII.

**Figure 2.**
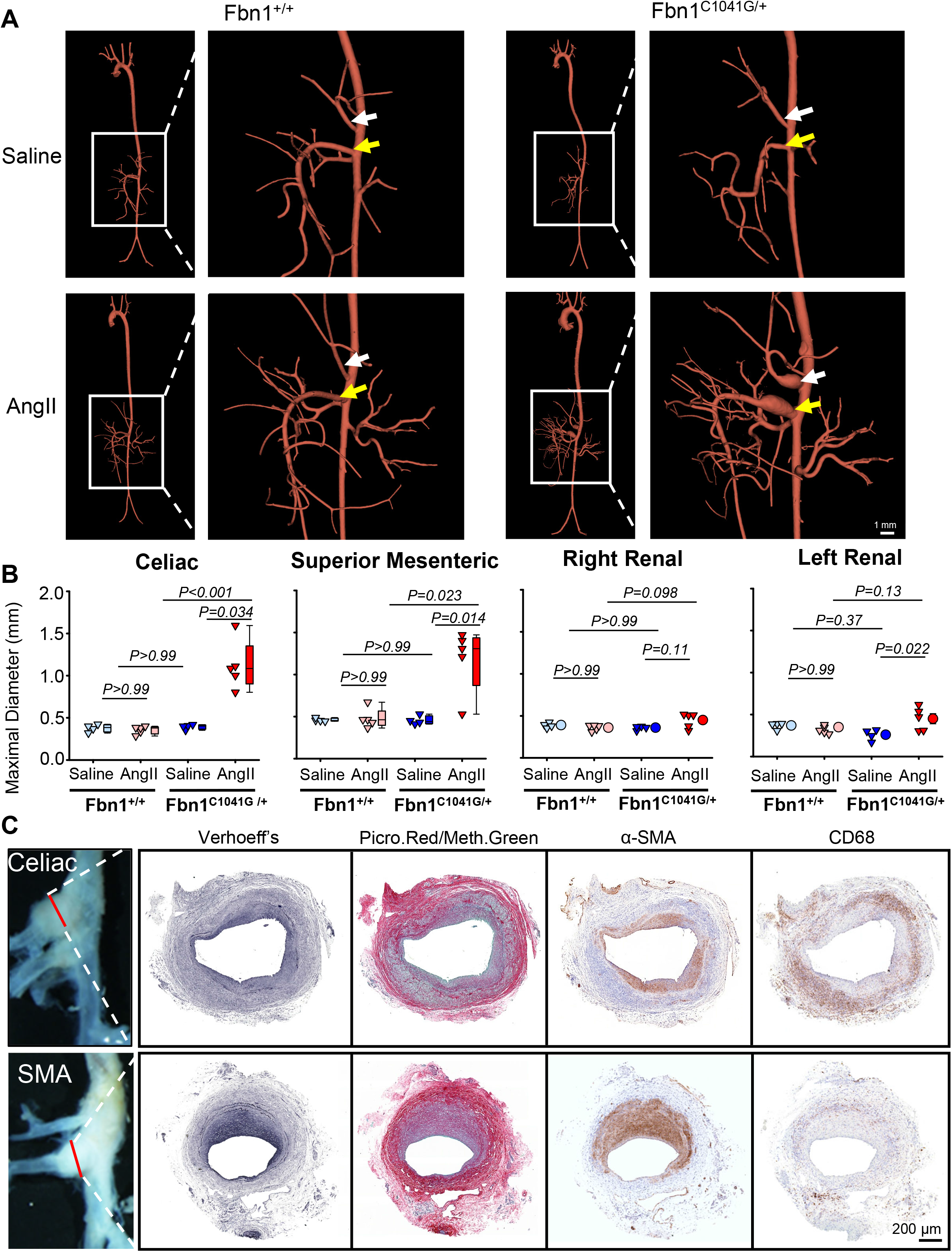
AngII Promoted Development of Aortic Branch Aneurysms in *Fbn1*^*C1041G/+*^ Mice. Nine- to 14-week-old male *Fbn1*^*+/+*^ and *Fbn1*^*C1041G/+*^ mice were infused with either saline or AngII for 28 days. (**A**) Representative microCT images of the entire aorta and the abdominal aorta. (**B**) Maximum diameters of the aortic branch points for celiac, superior mesenteric, right renal, and left renal arteries measured using microCT images. Data were analyzed using two-way ANOVA (nonparametric for celiac and superior mesenteric maximal diameters and parametric for right and left renal maximal diameters) followed by contrast tests with Bonferroni correction. N=4 in *Fbn1*^*+/+*^ saline, N=5 in *Fbn1*^*+/+*^AngII, N=4 in *Fbn1*^*C1041G/+*^ saline, and N=5 in *Fbn1*^*C1041G/+*^ AngII groups at termination. (**C**) Tissue sections of the celiac and superior mesenteric aortic branches were stained with Verhoeff’s iron hematoxylin, picrosirius red/methyl green, and immunostained for α-smooth muscle actin and CD68, respectively.

To determine whether the findings in AngII-infused mice were attributed to differences in blood pressure responses, we compared systolic blood pressure in *Fbn1*^*C1041G/+*^ mice infused with either AngII (1,000 ng/kg/min) or NE (5.6 mg/kg/day) in the same study. Because male *Fbn1*^*C1041G/+*^ mice exhibit high mortality following AngII infusion, AngII or NE was administered only to female *Fbn1*^*C1041G/+*^ mice for 28 days. Our data showed that systolic blood pressure did not differ between females infused with either AngII or NE (Figure S4A). Despite comparable blood pressures between the two infusion groups, pronounced aortic branch pathologies in the celiac and superior mesenteric arteries were observed only in AngII-infused mice, but not in NE-infused mice, as shown by microCT (Figure S4B). Consistently, the maximal diameters of these two aortic branches were larger in AngII-infused mice than in NE-infused mice (Figure S4C).

Given the presence of branch aneurysms in the suprarenal aortic region, ex vivo imaging did not allow reliable measurements of maximal aortic diameters in this area. To address this limitation, we measured maximal suprarenal aortic diameters using microCT images (Figure S6). AngII infusion modestly increased suprarenal aortic diameters in male *Fbn1*^*C1041G/+*^ mice, whereas no significant changes were observed in female *Fbn1*^*C1041G/+*^ mice or in male and female *Fbn1*^+/+^ mice. Subsequently, histological analysis of AngII-infused male *Fbn1*^*C1041G/+*^ mice was performed in regions encompassing the relatively preserved aortic segment above the celiac artery, the celiac branch point, and the superior mesenteric branch point (Figure S7-10). In any of these 3 locations, elastic lamellae of the aorta appeared largely intact, with restricted fragmentation. In contrast, marked elastic fiber disruption was observed within the walls of both arterial branches (Figure 7). These branch regions also exhibited pronounced adventitial collagen deposition (Figure S8), disrupted smooth muscle α-actin staining within the medial layers (Figure S9), and substantial accumulation of CD68-positive cells (Figure S10). Notably, the free wall of the adjacent aorta showed modest wall thickening with mild macrophage infiltration. These findings indicate that AngII-induced pathology in male *Fbn1*^*C1041G/+*^ mice is spatially restricted, with preferential and more severe involvement of the celiac and superior mesenteric arterial branch sites compared to the adjacent suprarenal aortic wall.

## DISCUSSION

There is an increasing awareness that *FBN1* variants can lead to a spectrum of aortic pathologies. For example, it has been recognized recently that branch aneurysms are frequent in Marfan Syndrome patients and are associated with vascular risk.^3^ This study has demonstrated that infusion of AngII into mice expressing the C1041G/+ variant of *Fbn1* leads to augmented aneurysms in the proximal thoracic aorta, accelerated arterial rupture, and aortic branch aneurysms.

Our previous studies demonstrated that the age-dependent expansion of the ascending aorta in *Fbn1*^C1041G/+^ mice has a strong sexual dimorphism, with males showing gradual dilation over the course of a year, whereas females display a rate of increase indistinguishable from that of *Fbn1*^+/+^ mice.^8^ These results were replicated in the current study. Sexual dimorphism was also present in some aspects of aortic diseases during AngII infusion. Specifically, there was a much greater incidence of death due to loss of aortic integrity in males compared to females. However, similar to males, female *Fbn1*^C1041G/+^ mice infused with AngII had striking expansion of the ascending aorta. This finding implies that distinct mechanisms underlie AngII-induced aneurysm formation and rupture in *Fbn1*^C1041G/+^ mice, which will need to be further elucidated.

A novel observation in this study is the presence of aortic branch aneurysms predominantly involving the celiac and superior mesenteric arteries in AngII-infused *Fbn1*^*C1041G/+*^ mice, irrespective of sex. This finding parallels recent clinical reports demonstrating aortic branch aneurysms in patients with Marfan syndrome.^3^ Consistent with these observations, the 2022 ACC/AHA guidelines recognize that aortic branch aneurysms are more common than previously appreciated, with an elevated likelihood of requiring aortic surgery.^26^ The mechanisms by which AngII preferentially promotes aneurysm formation in the celiac and superior mesenteric arteries of *Fbn1*^C1041G/+^ mice remain unclear. Both arteries supply blood to major gastrointestinal organs, and the superior mesenteric artery functions as a resistant artery critical for regulating regional blood flow and systemic blood pressure. Their unique hemodynamic demands, complex branching geometry, and distinct extracellular matrix architecture may render these two arteries susceptible to the combined effects of *Fbn1* haploinsufficiency and AngII-induced mechanical and inflammatory stress. Given their central roles in abdominal organ perfusion, further exploration is warranted to elucidate how AngII drives aneurysmal development and vascular remodeling in these specific arteries of *Fbn1*^C1041G/+^ mice.

Infusion of AngII at sufficiently high rates increases systolic blood pressure, which could potentially contribute to aortic branch pathology observed in this study. One approach to determine the contribution of blood pressure *per se* is to compare pathological outcomes in mice infused with either AngII or NE at rates that produce comparable increases in systolic blood pressure, as demonstrated in our previous studies.^20, 27-30^ In the present study, we used an NE infusion rate (5.6 mg/kg/day) that has been shown previously to increase systolic blood pressure similar to that produced by AngII infusion at 1,000 ng/kg/min. Despite producing a comparable blood pressure response, NE infusion failed to reproduce the aortic branch pathologies observed with AngII infusion. These findings imply that increased blood pressure alone is not sufficient to account for the development of these aortic branch pathologies and support the notion that AngII exerts blood pressure-independent effects that contribute to disease development.

In summary, this study demonstrates that AngII infusion in *Fbn1*^C1041G/+^ mice augments aneurysmal disease in the distal thoracic aorta and induces previously undocumented pathologies, including aortic rupture and aortic branch aneurysms. This model provides a valuable platform to perform subsequent studies to define the mechanisms of these diverse pathologies.

## Abbreviations

AngII: Angiotensin II
Fbn1: Fibrillin-1
microCT: Micro Computed Tomography
NE: Norepinephrine

## Acknowledgments

These studies were facilitated by the University of Kentucky Light Microscopy Core (RRID: SCR 026405) and the Magnetic Resonance and Spectroscopy Core (RRID: SCR 026383).

## REFERENCES

1. Milewicz DM, Braverman AC, De Backer J, Morris SA, Boileau C, Maumenee IH, Jondeau G, Evangelista A and Pyeritz RE. Marfan syndrome. Nat Rev Dis Primers. 2021;7:64.

2. Requejo-Garcia L, Martinez-Lopez R, Plana-Andani E, Medina-Badenes P, Hernandiz-Martinez A, Torres-Blanco A and Miralles-Hernandez M. Extrathoracic aneurysms in Marfan syndrome: A systematic review of the literature. Ann Vasc Surg. 2022;87:548–559.

3. Lopez-Sainz A, Mila L, Rodriguez-Palomares J, Limeres J, Granato C, La Mura L, Sabate A, Guala A, Gutierrez L, Galian-Gay L, Sao-Aviles A, Bellmunt S, Rodriguez R, Cuellar-Calabria H, Roque A, Ferreira-Gonzalez I, Evangelista A and Teixido-Tura G. Aortic branch aneurysms and vascular risk in patients with Marfan syndrome. J Am Coll Cardiol. 2021;77:3005–3012.

4. Summers KM. Genetic models of fibrillinopathies. Genetics. 2024;226.

5. Judge DP, Biery NJ, Keene DR, Geubtner J, Myers L, Huso DL, Sakai LY and Dietz HC. Evidence for a critical contribution of haploinsufficiency in the complex pathogenesis of Marfan syndrome. J Clin Invest. 2004;114:172–181.

6. Habashi JP, Doyle JJ, Holm TM, Aziz H, Schoenhoff F, Bedja D, Chen Y, Modiri AN, Judge DP and Dietz HC. Angiotensin II type 2 receptor signaling attenuates aortic aneurysm in mice through ERK antagonism. Science. 2011;332:361–365.

7. Gharraee N, Sun Y, Swisher JA and Lessner SM. Age and sex dependency of thoracic aortopathy in a mouse model of Marfan syndrome. Am J Physiol Heart Circ Physiol. 2022;322:H44–H56.

8. Chen JZ, Sawada H, Ye D, Katsumata Y, Kukida M, Ohno-Urabe S, Moorleghen JJ, Franklin MK, Howatt DA, Sheppard MB, Mullick AE, Lu HS and Daugherty A. Deletion of AT1a (Angiotensin II type 1a) receptor or inhibition of angiotensinogen synthesis attenuates thoracic aortopathies in fibrillin1(C1041G/+) mice. Arterioscler Thromb Vasc Biol. 2021;41:2538–2550.

9. Sawada H, Lu HS, Cassis LA and Daugherty A. Twenty years of studying AngII (angiotensin II)-induced abdominal aortic pathologies in mice: continuing questions and challenges to provide insight into the human disease. Arterioscler Thromb Vasc Biol. 2022;42:277–288.

10. Bush E, Maeda N, Kuziel WA, Dawson TC, Wilcox JN, DeLeon H and Taylor WR. CC chemokine receptor 2 is required for macrophage infiltration and vascular hypertrophy in angiotensin II-induced hypertension. Hypertension. 2000;36:360–363.

11. Daugherty A, Manning MW and Cassis LA. Angiotensin II promotes atherosclerotic lesions and aneurysms in apolipoprotein E-deficient mice. J Clin Invest. 2000;105:1605–1612.

12. Daugherty A and Cassis L. Chronic angiotensin II infusion promotes atherogenesis in low density lipoprotein receptor -/- mice. Ann NY Acad Sci. 1999;892:108–118.

13. Kanematsu Y, Kanematsu M, Kurihara C, Tsou TL, Nuki Y, Liang EI, Makino H and Hashimoto T. Pharmacologically induced thoracic and abdominal aortic aneurysms in mice. Hypertension. 2010;55:1267–1274.

14. Cavanaugh NB, Qian L, Westergaard NM, Kutschke WJ, Born EJ and Turek JW. A novel murine model of Marfan syndrome accelerates aortopathy and cardiomyopathy. Ann Thorac Surg. 2017;104:657–665.

15. Gensicke NM, Cavanaugh NB, Andersen ND, Huang T, Qian L, Dyle MC and Turek JW. Accelerated Marfan syndrome model recapitulates established signaling pathways. J Thorac Cardiovasc Surg. 2020;159:1719–1726.

16. Nettersheim FS, Lemties J, Braumann S, Geissen S, Bokredenghel S, Nies R, Hof A, Winkels H, Freeman BA, Klinke A, Rudolph V, Baldus S, Mehrkens D, Mollenhauer M and Adam M. Nitro-oleic acid reduces thoracic aortic aneurysm progression in a mouse model of Marfan syndrome. Cardiovasc Res. 2022;118:2211–2225.

17. Saddic L, Escopete S, Zilberberg L, Kalsow S, Gupta D, Eghbali M and Parker S. 17 beta-estradiol impedes aortic root dilation and rupture in male Marfan mice. Int J Mol Sci. 2023;24.

18. Roth L, Schrijvers DM, Martinet W and De Meyer GR. Angiotensin II increases coronary fibrosis, cardiac hypertrophy and the incidence of myocardial infarctions in ApoE(-/-)Fbn1(C1039G+/-) mice. Acta Cardiol. 2016;71:483–488.

19. Daugherty A, Milewicz DM, Dichek DA, Ghaghada KB, Humphrey JD, LeMaire SA, Li Y, Mallat Z, Saeys Y, Sawada H, Shen YH, Suzuki T, Zhou Z, Leducq Network on C and Molecular Drivers of Acute Aortic D. Recommendations for design, execution, and reporting of studies on experimental thoracic aortopathy in preclinical models. Arterioscler Thromb Vasc Biol. 2025;45:609–631.

20. Davis FM, Rateri DL, Balakrishnan A, Howatt DA, Strickland DK, Muratoglu SC, Haggerty CM, Fornwalt BK, Cassis LA and Daugherty A. Smooth muscle cell deletion of low-density lipoprotein receptor-related protein 1 augments angiotensin II-induced superior mesenteric arterial and ascending aortic aneurysms. Arterioscler Thromb Vasc Biol. 2015;35:155–162.

21. Sawada H, Katsumata Y, Higashi H, Zhang C, Li Y, Morgan S, Lee LH, Singh SA, Chen JZ, Franklin MK, Moorleghen JJ, Howatt DA, Rateri DL, Shen YH, LeMaire SA, Aikawa M, Majesky MW, Lu HS and Daugherty A. Second heart field-derived cells contribute to angiotensin II-mediated ascending aortopathies. Circulation. 2022;145:987–1001.

22. Daugherty A, Rateri D, Lu H and Balakrishnan A. Measuring blood pressure in mice using volume pressure recording, a tail-cuff method. J Vis Exp. 2009;27:1291.

23. Zhang JM, Au DT, Sawada H, Franklin MK, Moorleghen JJ, Howatt DA, Wang P, Aicher BO, Hampton B, Migliorini M, Ni F, Mullick AE, Wani MM, Ucuzian AA, Lu HS, Muratoglu SC, Daugherty A and Strickland DK. LRP1 protects against excessive superior mesenteric artery remodeling by modulating angiotensin II-mediated signaling. JCI Insight. 2023;8:e164751.

24. Ohno-Urabe S, Kukida M, Franklin MK, Katsumata Y, Su W, Gong MC, Lu HS, Daugherty A and Sawada H. Authentication of in situ measurements for thoracic aortic aneurysms in mice. Arterioscler Thromb Vasc Biol. 2021;41:2117–2119.

25. Li W, Li Q, Jiao Y, Qin L, Ali R, Zhou J, Ferruzzi J, Kim RW, Geirsson A, Dietz HC, Offermanns S, Humphrey JD and Tellides G. Tgfbr2 disruption in postnatal smooth muscle impairs aortic wall homeostasis. J Clin Invest. 2014;124:755–767.

26. Isselbacher EM, Preventza O, Hamilton Black J, 3rd, Augoustides JG, Beck AW, Bolen MA, Braverman AC, Bray BE, Brown-Zimmerman MM, Chen EP, Collins TJ, DeAnda A, Jr., Fanola CL, Girardi LN, Hicks CW, Hui DS, Schuyler Jones W, Kalahasti V, Kim KM, Milewicz DM, Oderich GS, Ogbechie L, Promes SB, Gyang Ross E, Schermerhorn ML, Singleton Times S, Tseng EE, Wang GJ and Woo YJ. 2022 ACC/AHA guideline for the diagnosis and management of aortic disease: A report of the American Heart Association/American College of Cardiology Joint Committee on Clinical Practice Guidelines. Circulation. 2022;146:e334–e482.

27. Owens APr, Subramanian V, Moorleghen JJ, Guo Z, McNamara CA, Cassis LA and Daugherty A. Angiotensin II induces a region-specific hyperplasia of the ascending aorta through regulation of inhibitor of differentiation 3. Circ Res. 2010;106:611–619.

28. Rateri DL, Davis FM, Balakrishnan A, Howatt DA, Moorleghen JJ, O’Connor WN, Charnigo R, Cassis LA and Daugherty A. Angiotensin II induces region-specific medial disruption during evolution of ascending aortic aneurysms. Am J Pathol. 2014;184:2586–2595.

29. Pettey AC, Ito S, Franklin MK, Howatt DA, Moorleghen JJ, Levitan BM, Graf DB, Guzman VZ, Zhang N, Sawada H, Saffitz JE, Lu HS and Daugherty A. Cardiac hemorrhage precedes hypertension-induced fibrosis in plasminogen activator inhibitor-1 deficient mice. bioRxiv. 2025: doi: 10.1101/2025.11.19.689269.

30. Cassis LA, Gupte M, Thayer S, Zhang X, Charnigo R, Howatt DA, Rateri DL and Daugherty A. ANG II infusion promotes abdominal aortic aneurysms independent of increased blood pressure in hypercholesterolemic mice. Am J Physiol Heart Circ Physiol. 2009;296:H1660–1665.

