## Supplementary material for "Angiotensin II Induces Abdominal Aortic Branch Aneurysms in *Fibrillin-1^C1041G/+^* Mice": Tables and Figures

[1] Saha Cardiovascular Research Center
[2] Saha Aortic Center
[3] Department of Physiology,
[4] Department of Family and Community Medicine
[5] Sanders-Brown Center on Aging,
[6] Division of Biomedical Informatics
University of Kentucky, Lexington, KY

Running title – Aortic Branch Aneurysms

**Corresponding Authors:**
Alan Daugherty
Saha CVRC, BBSRB, Room B243
University of Kentucky
741 S Limestone
Lexington KY 40356-0509

Hong S. Lu
Saha CVRC, BBSRB, Room B249
University of Kentucky
741 S Limestone
Lexington KY 40356-0509

**Table S1. Necropsy on *Fbn1*$^{C1041G/+}$ mice that died during AngII infusion**

| **Sex** | **From Initiation of AngII Infusion to Death** | **Cause of Death (Number of Mice)** | | |
|---|---|---|---|---|
| | Time (Days) | Blood Clot Found in Thoracic Cavity | Blood Clot Found in Abdominal Cavity | Nonvascular-based Death |
| **Male** | 2 | N=2 | N=2 | |
| | 3 | | N=4 | |
| | 4 | N=1 | | |
| | 5 | | N=1 | |
| | 6 | N=1 | | |
| | 15 | N=1 | | |
| | 16 | N=1 | | |
| | 19 | | | N=2 |
| | 22 | N=1 | | |
| | 25 | | | N=1 |
| | **Total** | **N=7** | **N=7** | **N=3** |
| **Female** | 4 | N=1 | | |
| | 8 | N=2 | | |
| | 26 | | | N=1 |
| | **Total** | N=3 | N=0 | N=1 |

### MAJOR RESOURCES TABLES

#### Table I. Mouse Models Used in This Study

| Strain | Vendor | Background Strain | Sex | Other Information | Persistent ID/URL |
|---|---|---|---|---|---|
| *Fbn1*$^{C1041G/+}$<br>*Fbn1*$^{+/+}$ | JAX (Strain # 012885) | C57BL/6J | Male and Female | Breeding Scheme:<br>Male *Fbn1*$^{C1041G/+}$ x Female *Fbn1*$^{+/+}$<br>Offspring:<br>*Fbn1*$^{+/+}$ and *Fbn1*$^{C1041G/+}$ littermates | https://www.jax.org/strain/012885 |

#### Table II. Primary Antibodies for Immunostaining

| Antibody | Vendor | Cat # | Working Concentration |
|---|---|---|---|
| Rabbit anti-smooth muscle α-actin | abcam | ab5694 | 2 μg/mL |
| Rabbit anti-CD68 (E3O7V) | Cell Signaling Technology | 97778 | 0.1 μg/mL |
| Rabbit nonimmune IgG | ImmunoReagents | Rb-003-V | Same as the antibody of interest (0.1-2 μg/mL, depending on primary antibodies) |

#### Table III. Secondary Antibodies for Immunostaining

| Antibody | Vendor | Cat # | Note |
|---|---|---|---|
| ImmPress Goat Anti-Rabbit IgG | Vector | MP-7451 | No concentration information available; Ready-to-use (per manufacturer) |

#### Table IV. Animal Study Information Following the ARRIVE Essential 10

##### Figure 1A, F, and K

| Groups | Sex | Age (weeks)* | Number (Start) | Number (Termination) |
|---|---|---|---|---|
| *Fbn1*$^{+/+}$ Saline | Male | 9-14 | 7 | 7 |
| *Fbn1*$^{+/+}$ AngII | Male | 9-14 | 20 | 20 |
| *Fbn1*$^{C1041G/+}$ Saline | Male | 9-14 | 7 | 6 |
| *Fbn1*$^{C1041G/+}$ AngII | Male | 9-14 | 26 | 9 |

##### Figure 1B

| Groups | Sex | Age (weeks)* | Thoracic Aortic Rupture | Abdominal Aortic Rupture | Non-Vascular |
|---|---|---|---|---|---|
| *Fbn1*$^{C1041G/+}$ AngII | Male | 9-14 | 7 | 7 | 3 |

##### Figure 1C, H, and L

| Groups | Sex | Age (weeks)* | Number (Start) | Number (Termination) |
|---|---|---|---|---|
| *Fbn1*$^{+/+}$ Saline | Female | 9-14 | 4 | 4 |
| *Fbn1*$^{+/+}$ AngII | Female | 9-14 | 22 | 21 |
| *Fbn1*$^{C1041G/+}$ Saline | Female | 9-14 | 4 | 4 |
| *Fbn1*$^{C1041G/+}$ AngII | Female | 9-14 | 20 | 16 |

**Figure 1D**

| Groups | Sex | Age (weeks)* | Thoracic Aortic Rupture | Abdominal Aortic Rupture | Non-Vascular |
|---|---|---|---|---|---|
| *Fbn1*$^{C1041G/+}$ AngII | Female | 9-14 | 3 | 0 | 1 |

**Figure 2B**

| Groups | Sex | Age (weeks)* | Number |
|---|---|---|---|
| *Fbn1*$^{+/+}$ Saline | Male | 9-14 | 4 |
| *Fbn1*$^{+/+}$ AngII | Male | 9-14 | 5 |
| *Fbn1*$^{C1041G/+}$ Saline | Male | 9-14 | 4 |
| *Fbn1*$^{C1041G/+}$ AngII | Male | 9-14 | 5 |

**Figure S2A, C, F, and J**

| Groups | Sex | Age (weeks)* | Number (Start) | Number (Termination) |
|---|---|---|---|---|
| *Fbn1*$^{+/+}$ Vehicle | Male | 9-14 | 4 | 4 |
| *Fbn1*$^{+/+}$ NE | Male | 9-14 | 4 | 4 |
| *Fbn1*$^{C1041G/+}$ Vehicle | Male | 9-14 | 5 | 5 |
| *Fbn1*$^{C1041G/+}$ NE | Male | 9-14 | 5 | 5 |

**Figure S2B, D, H, and L**

| Groups | Sex | Age (weeks)* | Number (Start) | Number (Termination) |
|---|---|---|---|---|
| *Fbn1*$^{+/+}$ Vehicle | Female | 9-14 | 5 | 5 |
| *Fbn1*$^{+/+}$ NE | Female | 9-14 | 5 | 5 |
| *Fbn1*$^{C1041G/+}$ Vehicle | Female | 9-14 | 6 | 6 |
| *Fbn1*$^{C1041G/+}$ NE | Female | 9-14 | 6 | 6 |

**Figure S3A**

| Groups | Sex | Age (weeks)* | Number |
|---|---|---|---|
| *Fbn1*$^{+/+}$ Saline | Female | 9-14 | 4 |
| *Fbn1*$^{+/+}$ AngII | Female | 9-14 | 4 |
| *Fbn1*$^{C1041G/+}$ Saline | Female | 9-14 | 4 |
| *Fbn1*$^{C1041G/+}$ AngII | Female | 9-14 | 7 |

**Figure S4A**

| Groups | Sex | Age (weeks)* | Number (Start) | Number (Termination) |
|---|---|---|---|---|
| *Fbn1*$^{C1041G/+}$ AngII | Female | 28 | 6 | 6 |
| *Fbn1*$^{C1041G/+}$ NE | Female | 28 | 6 | 6 |

**Figure S4B**

| Groups | Sex | Age (weeks)* | Number (Start) | Number (Termination) |
|---|---|---|---|---|
| *Fbn1*$^{C1041G/+}$ AngII | Female | 28 | 6 | 5 |
| *Fbn1*$^{C1041G/+}$ NE | Female | 28 | 6 | 4 |

##### Figure S6B - Male

| Groups | Sex | Age (weeks)* | Number |
|---|---|---|---|
| *Fbn1*$^{+/+}$ Saline | Male | 9-14 | 4 |
| *Fbn1*$^{+/+}$ AngII | Male | 9-14 | 5 |
| *Fbn1*$^{C1041G/+}$ Saline | Male | 9-14 | 4 |
| *Fbn1*$^{C1041G/+}$ AngII | Male | 9-14 | 5 |

##### Figure S6B - Female

| Groups | Sex | Age (weeks)* | Number |
|---|---|---|---|
| *Fbn1*$^{+/+}$ Saline | Female | 9-14 | 4 |
| *Fbn1*$^{+/+}$ AngII | Female | 9-14 | 4 |
| *Fbn1*$^{C1041G/+}$ Saline | Female | 9-14 | 4 |
| *Fbn1*$^{C1041G/+}$ AngII | Female | 9-14 | 7 |

Age (weeks)*: the age when AngII, NE, or their corresponding vehicle administration was started.

##### ARRIVE Essential 10 Checklist

| Item | Application |
|---|---|
| Ethics | Approved by the University of Kentucky IACUC (Protocol #: 2018-2967). |
| Sex | Described in each figure. |
| Inclusion criteria | Based on sex, age, body weight, and overall health appearance in each experiment. |
| Exclusion criteria | Based on sex, age, body weight, or medical cases reported by a veterinarian. |
| Sample size | Described in Major Resources Tables, Table IV. |
| Sample size calculation | None |
| Primary endpoint | Aortic and aortic branch pathologies |
| Randomization | Study mice were numbered and grouped randomly based on their genotype and sex. |
| Blinding | Maximal arterial diameters were measured by an investigator blinded to the study group information. Some data were measured by two investigators independently to validate the consistency of measurements. |
| Statistical analysis | SigmaPlot 16.0 (SYSTAT Software Inc., CA) or R Statistical Software. |
| Statistical method | Described in each figure legend. |
| Data availability | All numerical data used for figures and supplemental figures are available in the Supplemental Excel File. |

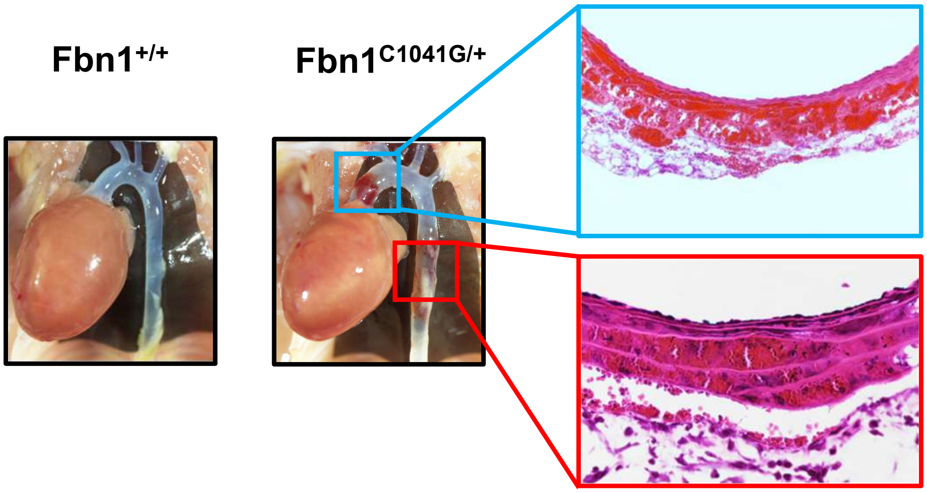

**Figure S1. AngII Infusion Led to Rapid Dissection of the Thoracic Aorta in *Fbn1$^{C1041G/+}$* Mice.** In situ images of the ascending and descending thoracic aorta of male *Fbn1$^{+/+}$* and *Fbn1$^{C1041G/+}$* mice infused with AngII for 3 days.

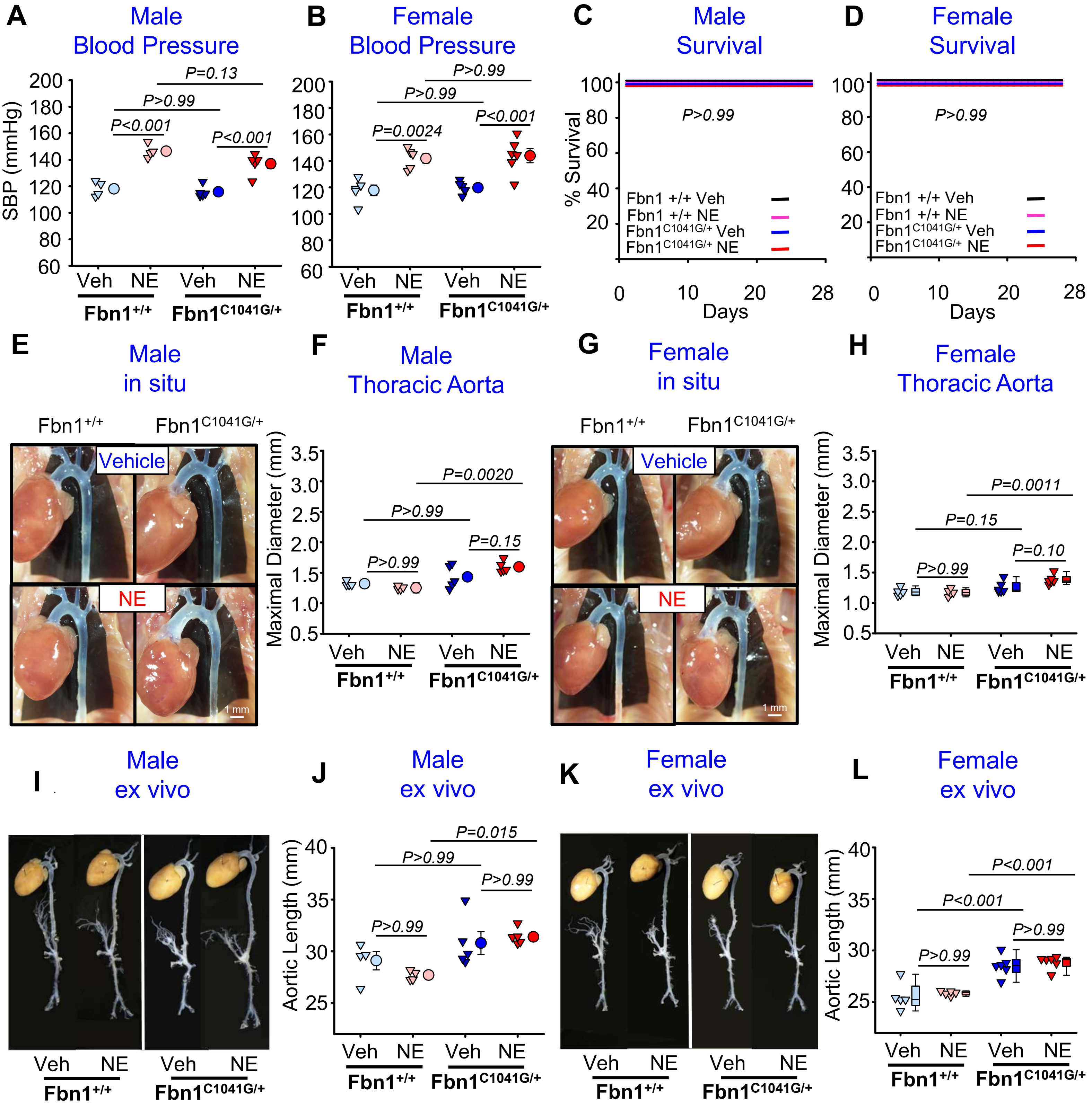

**Figure S2. NE Promoted Minimal Aortic Pathologies in *Fbn1*$^{+/+}$ and *Fbn1*$^{C1041G/+}$ Mice.** Nine- to 14-week-old male and female *Fbn1*$^{+/+}$ and *Fbn1*$^{C1041G/+}$ mice were infused with either vehicle (Veh) or NE for 28 days. Systolic blood pressure measurements in male (**A**) and female (**B**) mice. Survival curves for male (**C**) and female (**D**) mice were analyzed using a log-rank test. Representative in situ images of the thoracic aorta and maximal diameters of ascending aortas in male (**E-F**) and female (**G-H**) mice. Representative ex vivo images of the entire aorta in male (**I**) and female (**K**) mice. Aortic length measured from the left subclavian branch to the iliac bifurcation in male (**J**) and female (**L**) mice. Data were analyzed using two-way ANOVA (parametric in **A**, **B**, **F** and **J** and nonparametric in **H** and **L**) followed by contrast tests with Bonferroni correction. Animal numbers are presented in Table IV of the Major Resources Tables.

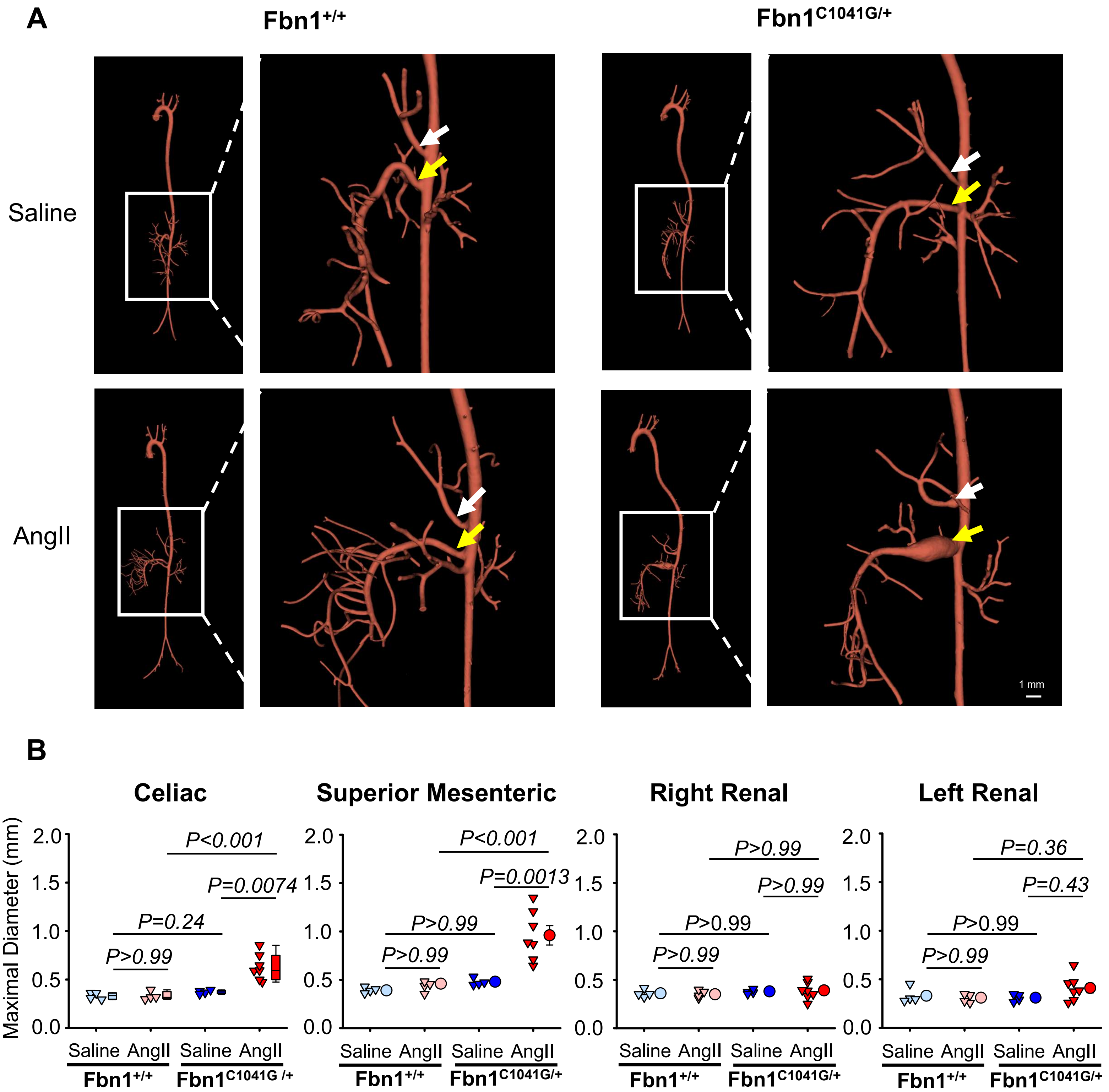

**Figure S3. AngII Infusion Promoted Development of Abdominal Aortic Brach Aneurysms in Female *Fbn1$^{C1041G/+}$* Mice.** Nine- to 14-week-old female *Fbn1$^{+/+}$* and *Fbn1$^{C1041G/+}$* mice were infused with either saline or AngII for 28 days. (**A**) Representative microCT images of the entire aorta and the abdominal aorta. (**B**) Maximum diameters of the aortic branch points for celiac, superior mesenteric, right renal, and left renal arteries measured using microCT images. Statistical analysis: two-way ANOVA (nonparametric for celiac artery and parametric for all the other arteries) followed by contrast tests with Bonferroni correction. Animal numbers are presented in Table IV of the Major Resources Tables.

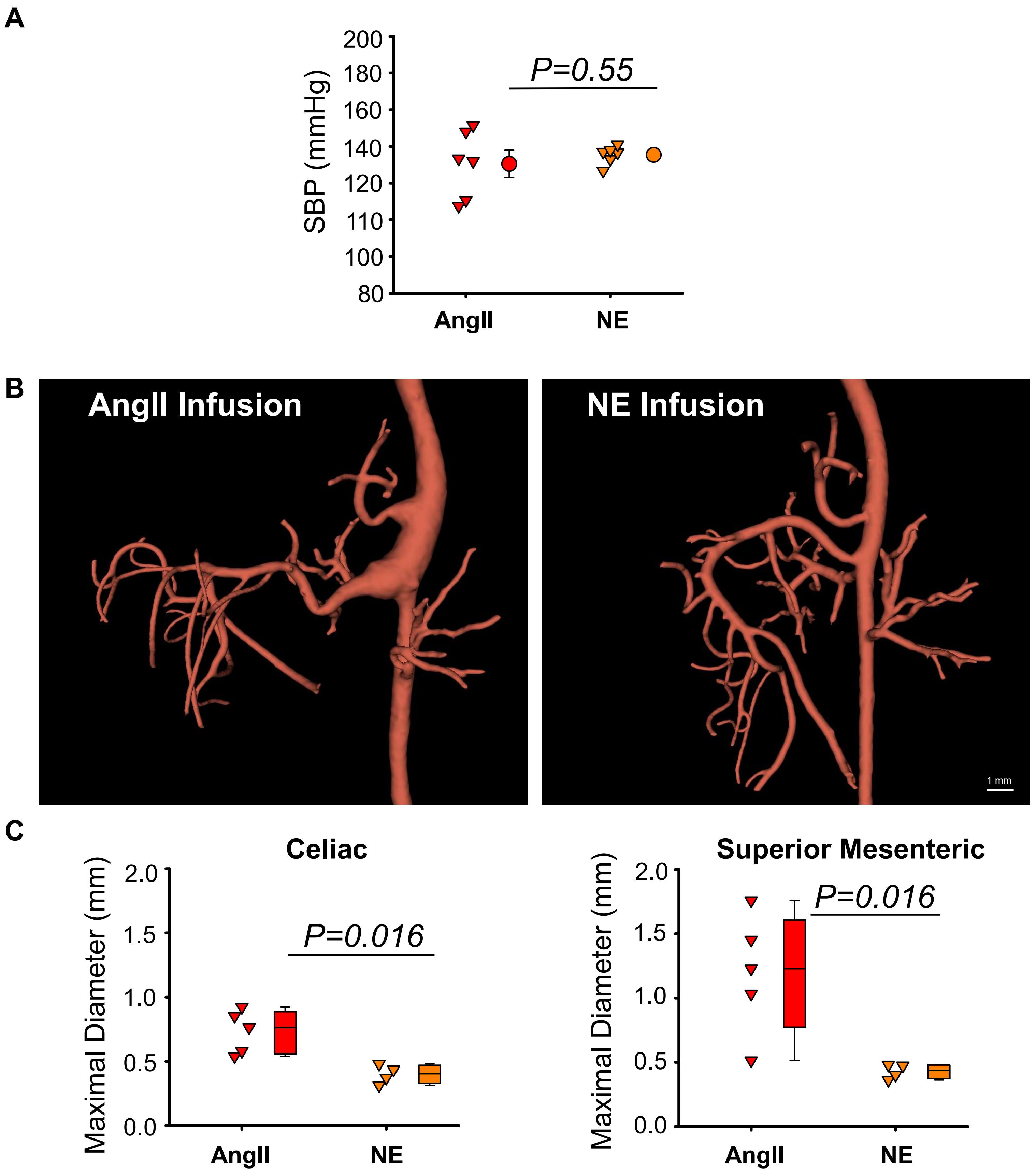

**Figure S4. AngII, but not NE, Promoted Aortic Branch Pathologies in Female *Fbn1*$^{C1041G/+}$ Mice.** Seven-month-old female *Fbn1*$^{C1041G/+}$ mice were infused with either AngII (1,000 ng/kg/min) or NE (5.6 mg/kg/day) for 28 days. (**A**) Systolic blood pressure measurements by a tail-cuff system. Statistical analysis: Welch's t-test. N=6/group. (**B**) Representative micro-CT images. (C) Maximal diameters of celiac and superior mesenteric arteries measured using microCT images. Statistical analysis: Mann-Whitney Rank Sum test. N=5 in AngII-infused mice and N=4 in NE-infused mice.

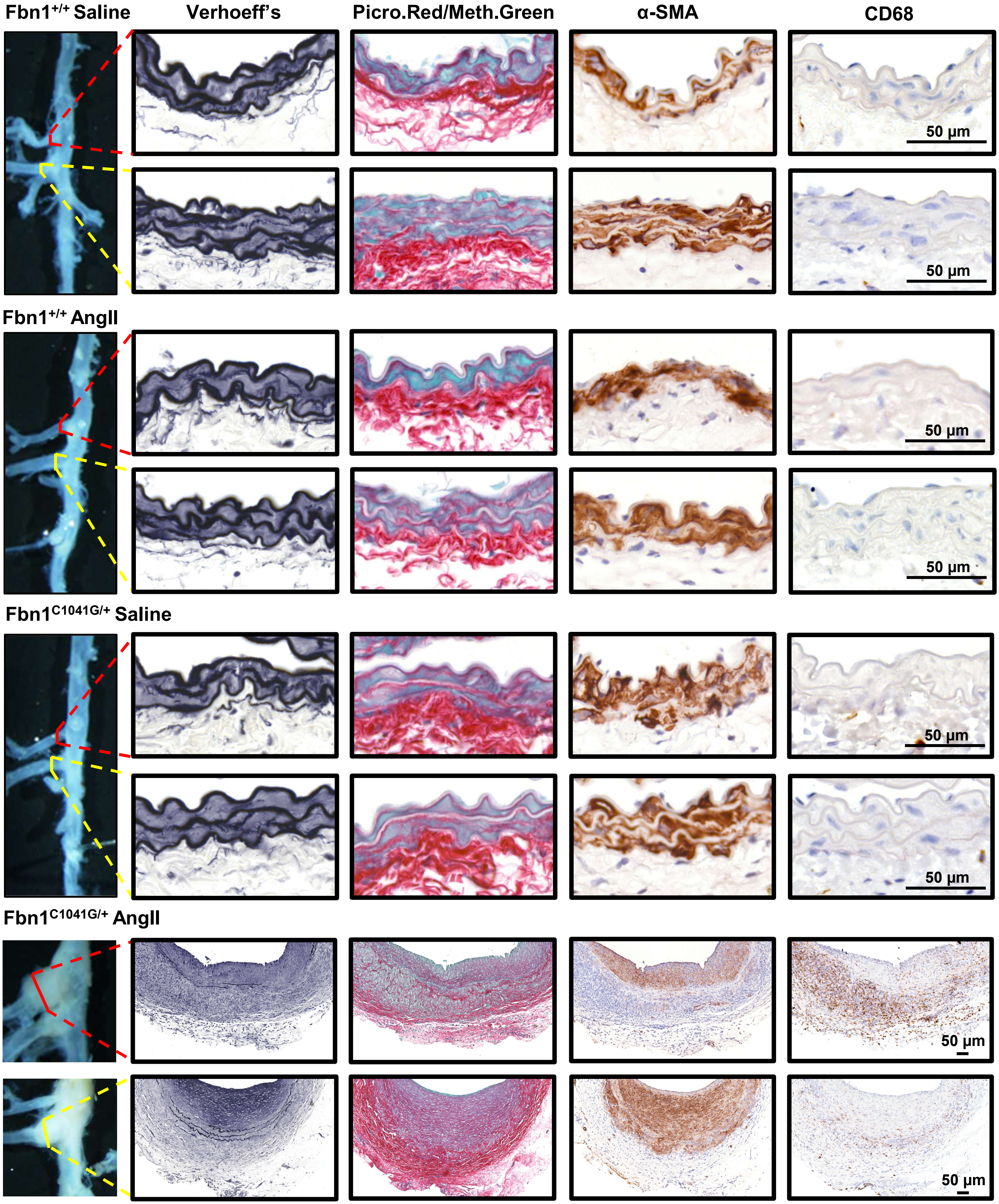

**Figure S5. Pathological Characterization of the Celiac and Superior Mesenteric Arteries in male mice.** Nine- to 14-week-old male *Fbn1*$^{+/+}$ and *Fbn1*$^{C1041G/+}$ mice were infused with either saline or AngII for 28 days. Arterial tissue sections were stained with Verhoeff's iron hematoxylin, picrosirius red/methyl green, and immunostained for α-smooth muscle actin and CD68, respectively.

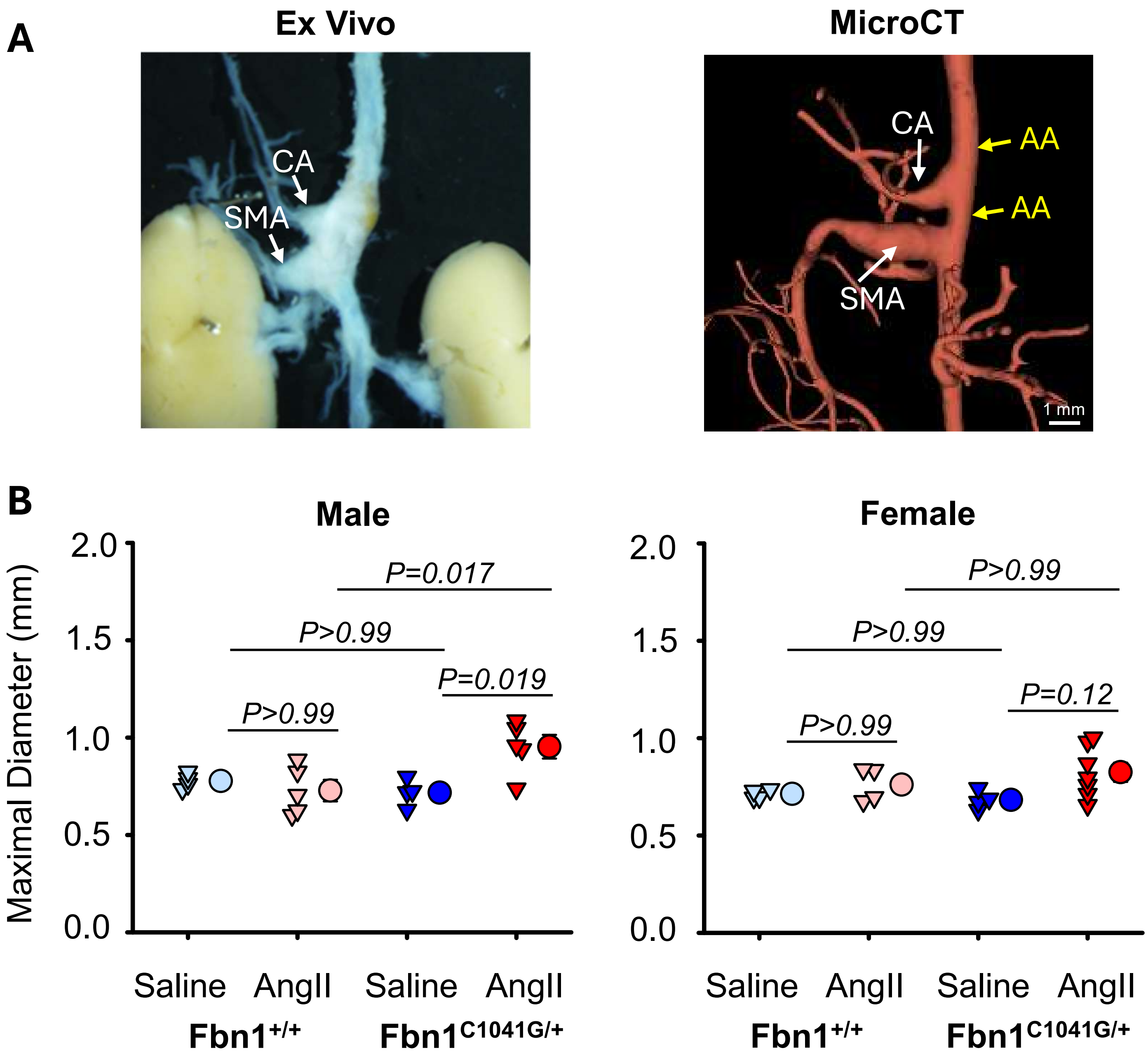

**Figure S6. AngII Infusion Led to Modest Increases of Suprarenal Abdominal Aortic Diameters in Male *Fbn1$^{C1041G/+}$* Mice.** Nine- to 14-week-old male and female *Fbn1$^{+/+}$* and *Fbn1$^{C1041G/+}$* mice were infused with either saline or AngII for 28 days. (**A**). Example of Ex Vivo image and microCT images. (**B**) microCT images were used to measure maximal diameters of the suprarenal aortic region. Statistical analysis: two-way ANOVA (parametric) followed by contrast tests with Bonferroni correction. Animal numbers are presented in Table IV of the Major Resources Tables. CA: celiac artery; SMA: superior mesenteric artery; AA: abdominal aorta

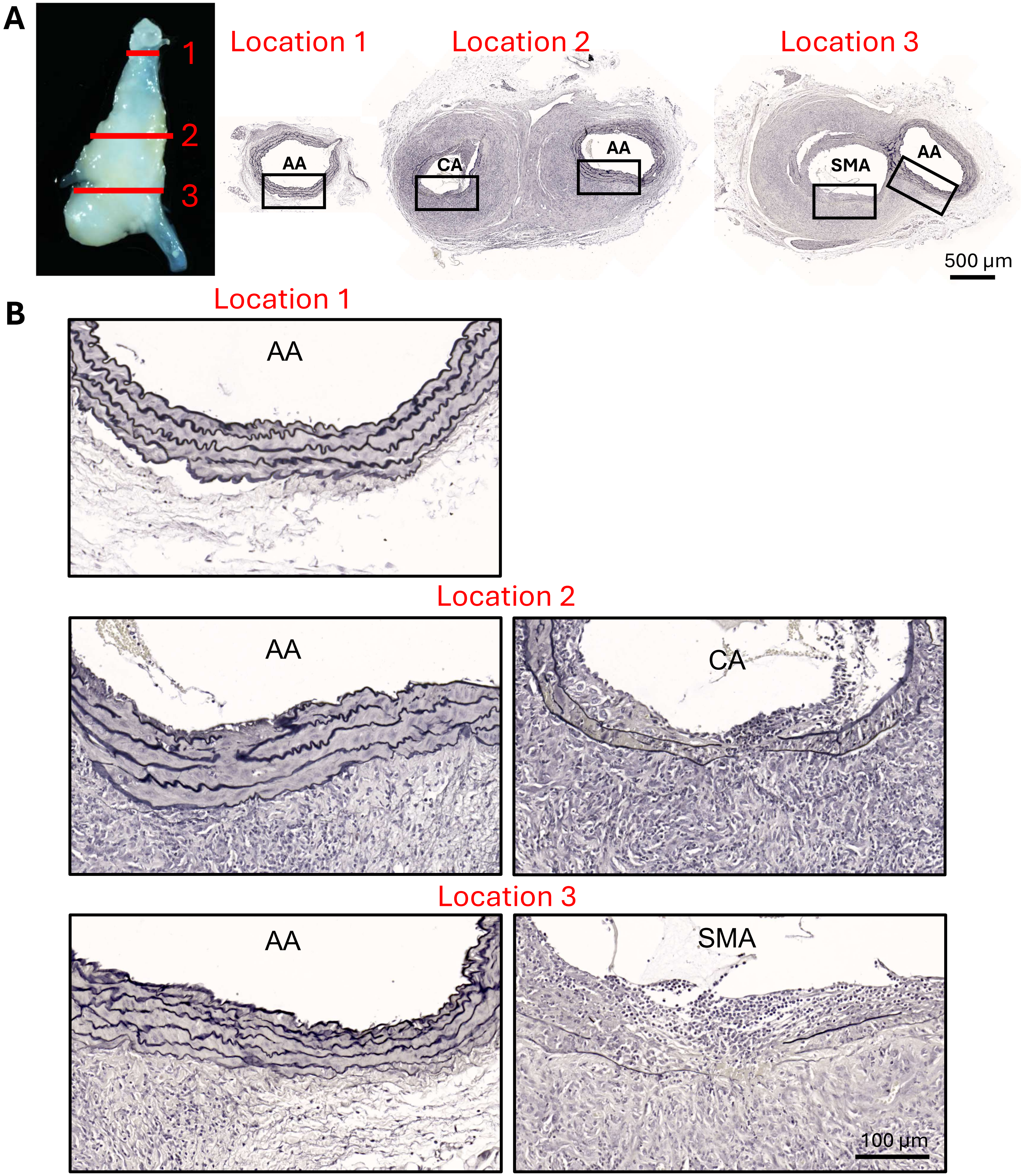

**Figure S7. AngII Infusion Led to Elastic Fiber Fragmentation Surrounding the Celiac and Superior Mesenteric Arteries.** Ex vivo image (**A**) is from a nine-week-old male *Fbn1*$^{C1041G/+}$ mouse infused with AngII for 28 days. Locations 1, 2, and 3 denote the anatomical sites where cross-sections were collected. Verhoff staining is shown at lower (**A**) and higher (**B**) magnifications. AA: abdominal aorta, CA: celiac artery, SMA: superior mesenteric artery

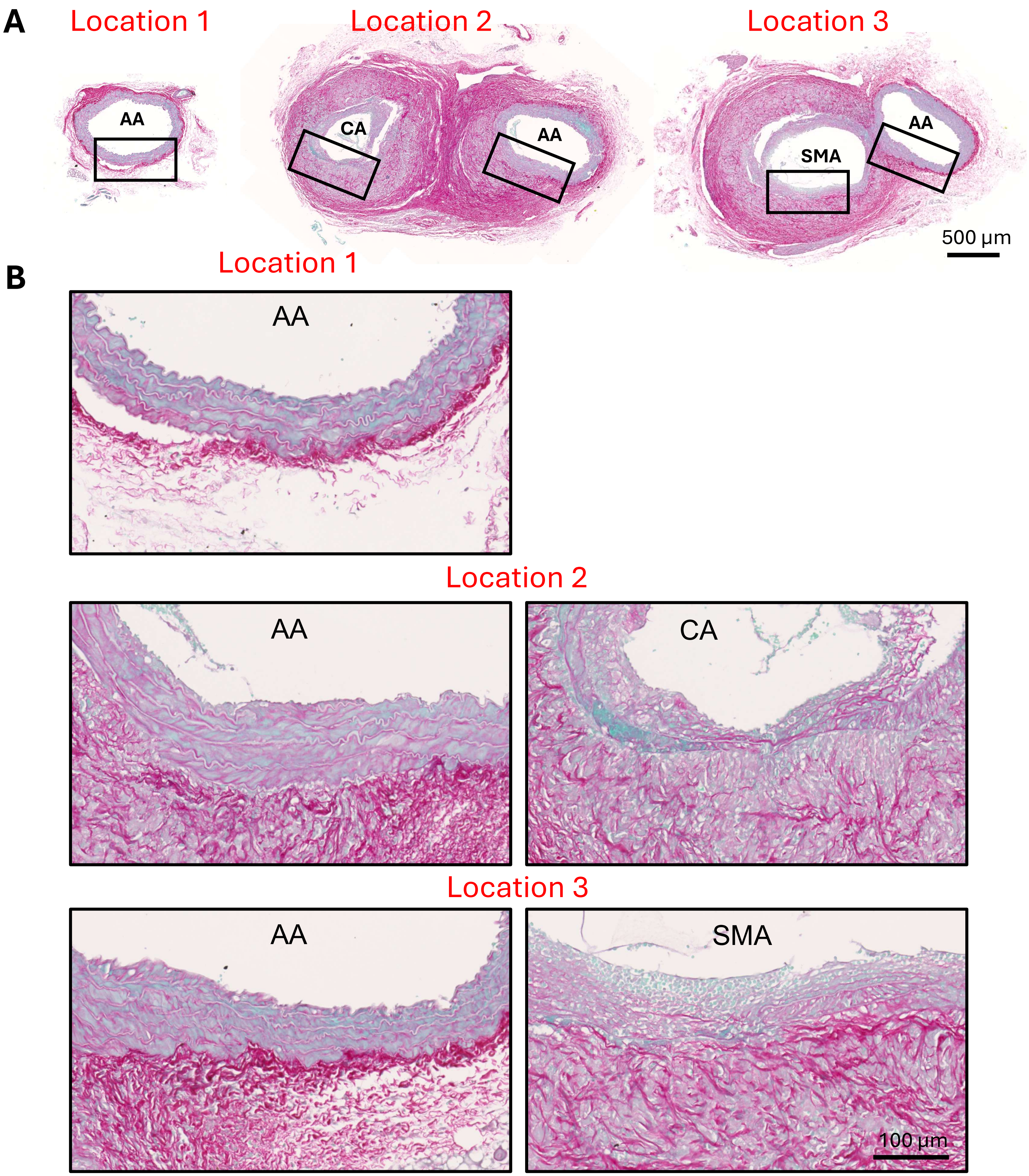

**Figure S8. AngII Infusion Led to Collagen Deposition Surrounding the Celiac and Superior Mesenteric Arteries.** Cross-sections were obtained from the same arterial tissues shown in Figure S7A. Locations 1, 2, and 3 denote the anatomical sites where cross-sections were collected. Picrosirius red and methyl green staining are shown at lower (**A**) and higher (**B**) magnifications. AA: abdominal aorta, CA: celiac artery, SMA: superior mesenteric artery.

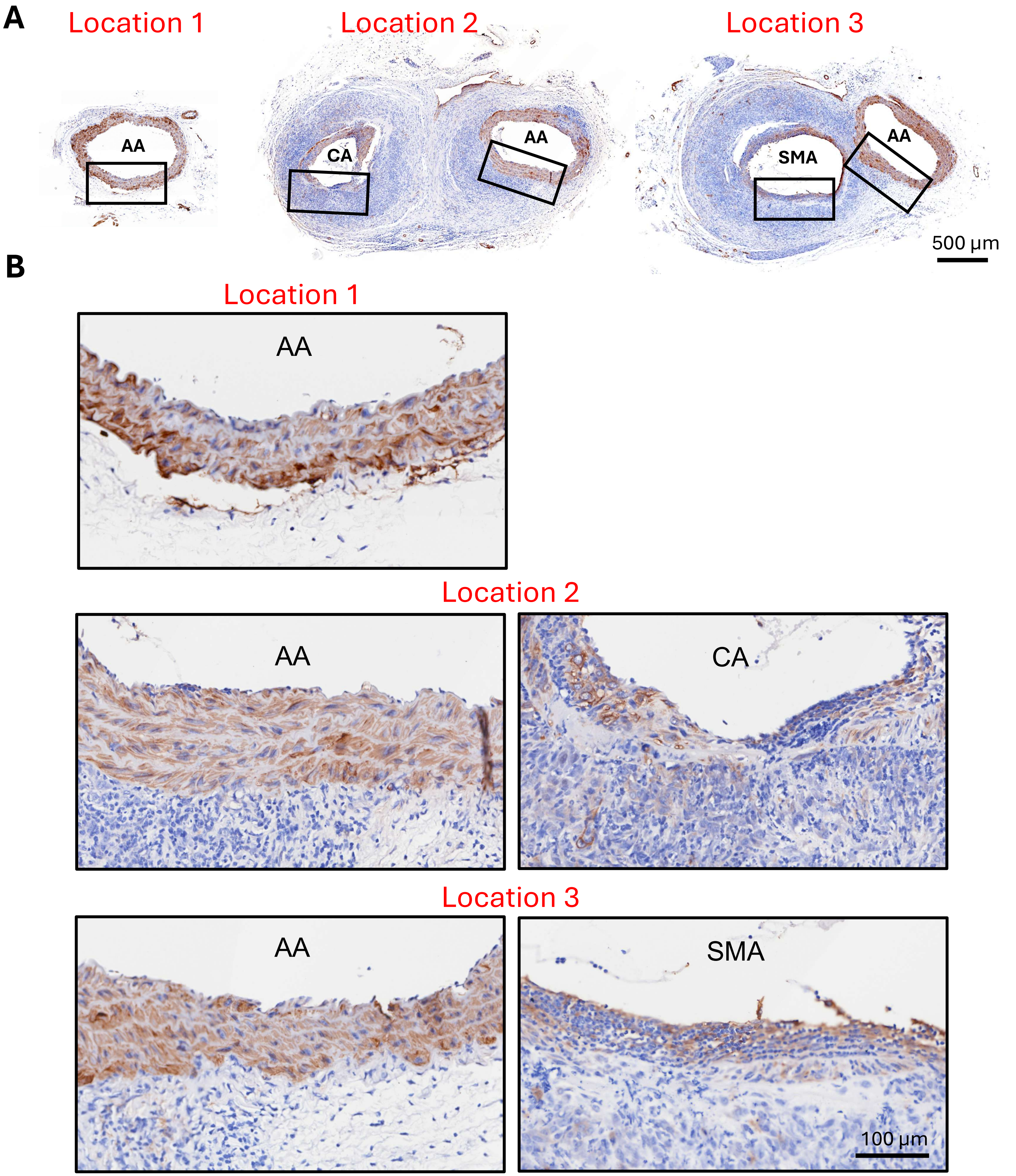

**Figure S9. AngII Infusion Led to Changes in Smooth Muscle Cells Surrounding the Celiac and Superior Mesenteric Arteries.** Cross-sections were obtained from the same arterial tissues shown in Figure S7A. Locations 1, 2, and 3 denote the anatomical sites where cross-sections were collected. Immunostaining of smooth muscle α-actin is shown at lower (**A**) and higher (**B**) magnifications. AA: abdominal aorta, CA: celiac artery, SMA: superior mesenteric artery.

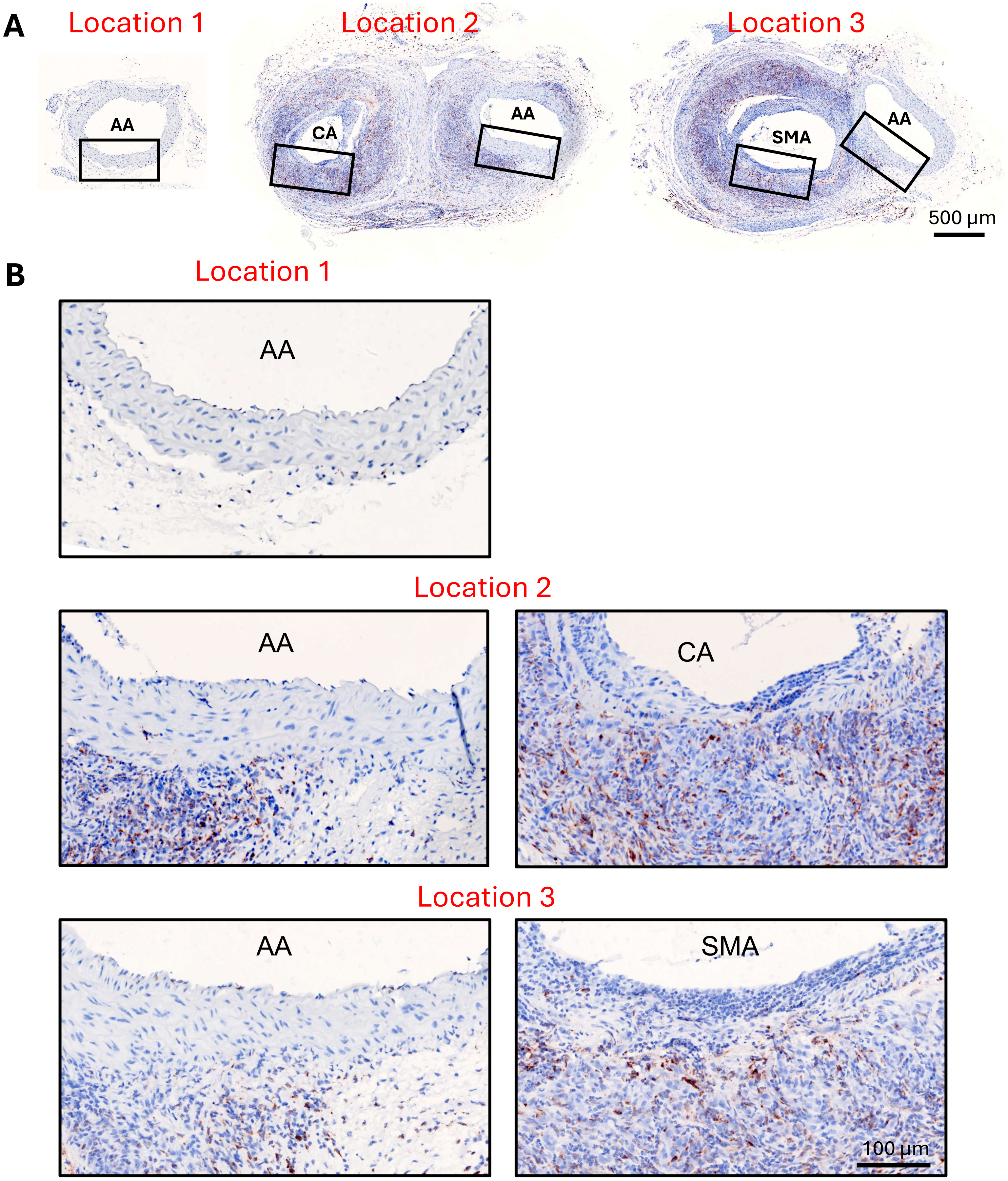

**Figure S10. AngII Infusion Led to Macrophage Accumulation Surrounding the Celiac and Superior Mesenteric Arteries.** Cross-sections were obtained from the same arterial tissues shown in Figure S7A. Locations 1, 2, and 3 denote the anatomical sites where cross-sections were collected. Immunostaining of CD68 is shown at lower (**A**) and higher (**B**) magnifications. AA: abdominal aorta, CA: celiac artery, SMA: superior mesenteric artery.
