## Supplementary figures and images for "Angiotensin II Induces Abdominal Aortic Branch Aneurysms in *Fibrillin-1^C1041G/+^* Mice"

### Videos

#### Slide 1
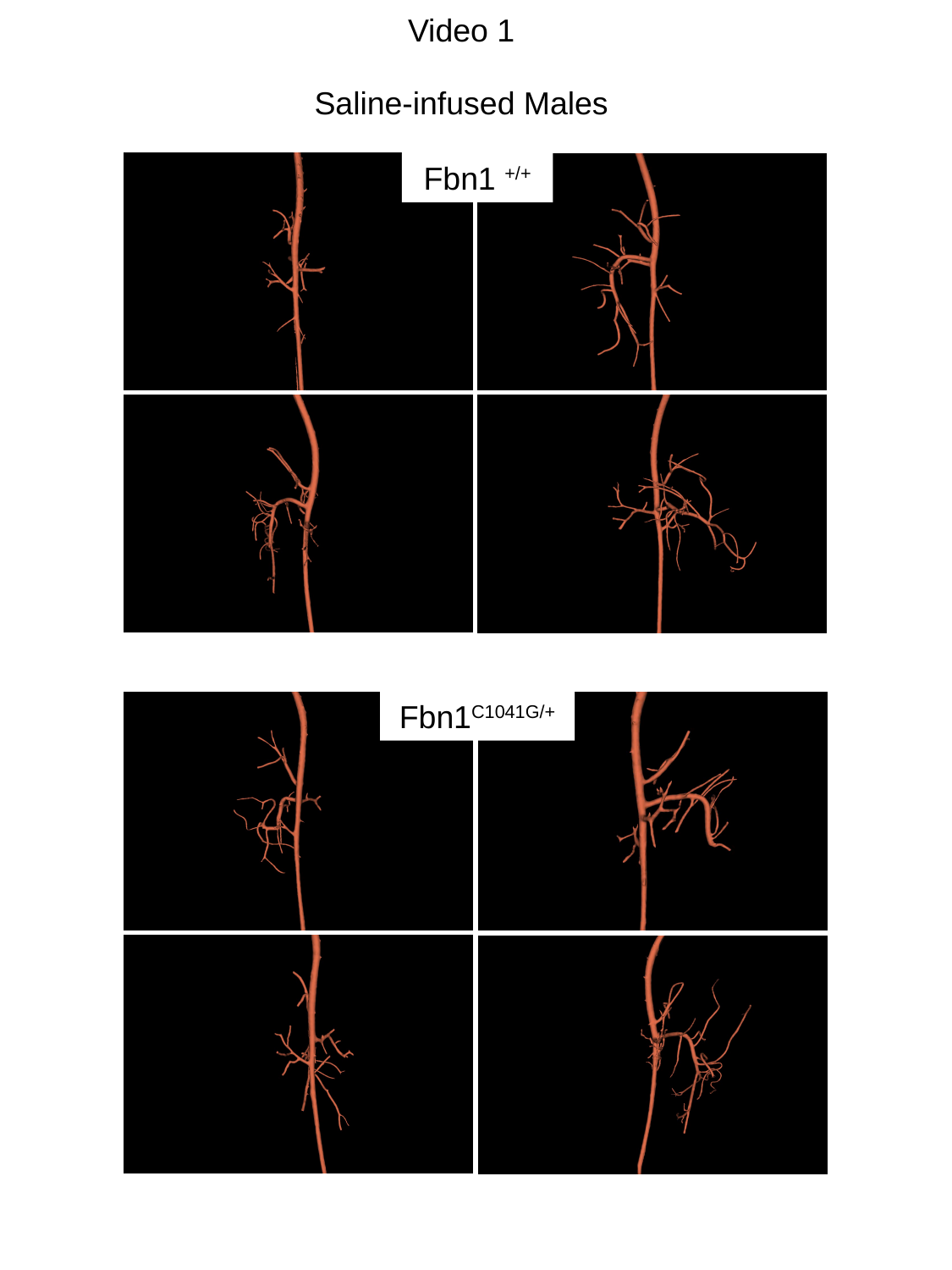

Video 1
Saline-infused Males
Fbn1 ^+/+^
Fbn1^C1041G/+^

#### Slide 2
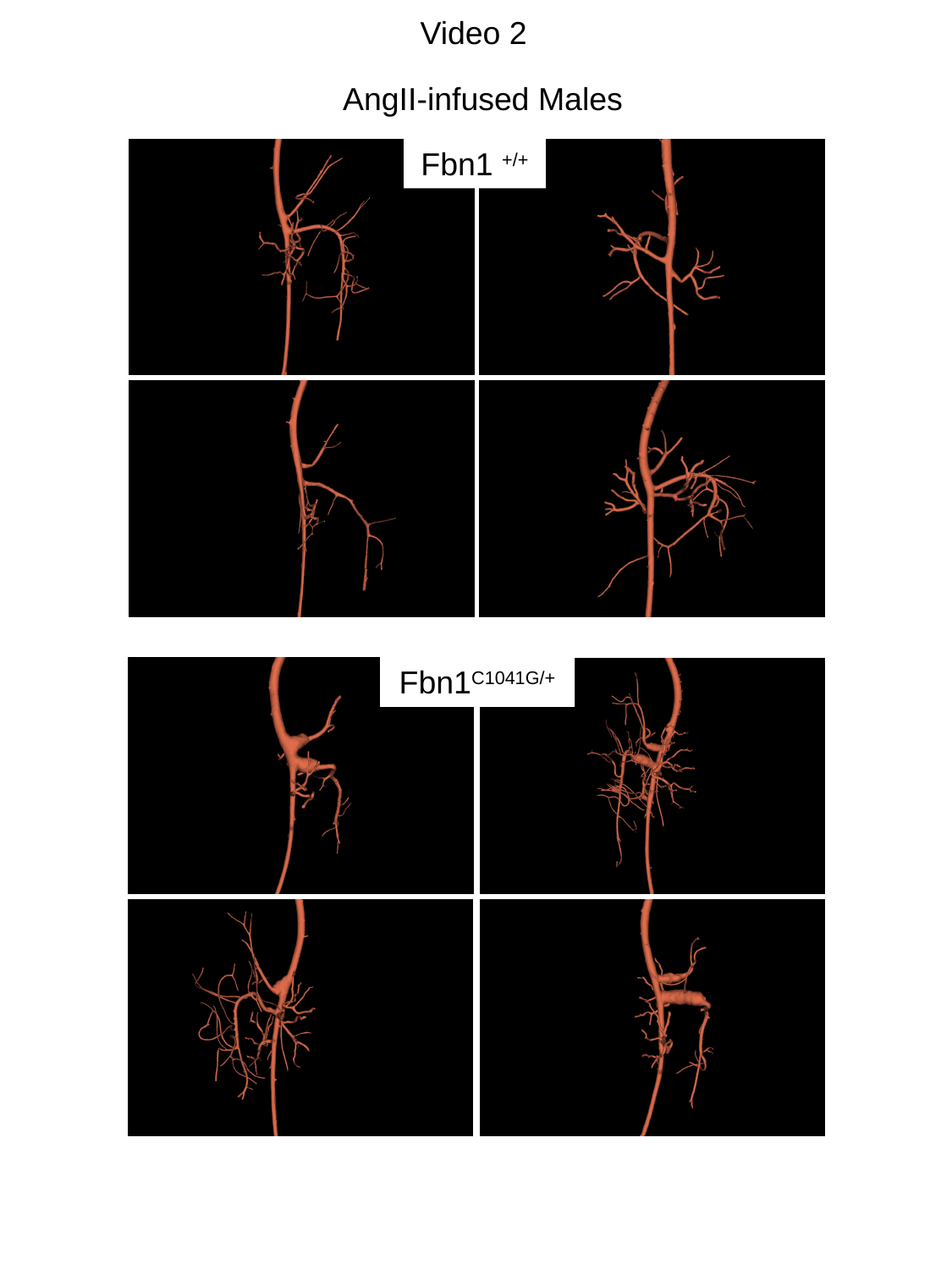

Video 2
AngII-infused Males
Fbn1 ^+/+^
Fbn1^C1041G/+^
